# Energy Expenditure during Cell Spreading Regulates the Stem Cells Responses to Matrix Stiffness

**DOI:** 10.1101/852400

**Authors:** Jing Xie, Min Bao, Xinyu Hu, Werner J. H. Koopman, Wilhelm T. S. Huck

**Author notes:** These authors contributed equally to this work (Jing Xie, Min Bao).

## Abstract

Cells respond to the mechanical properties of the extracellular matrix (ECM) through formation of focal adhesions (FAs), re-organization of the actin cytoskeleton and adjustment of cell contractility. These are energy-demanding processes, but a potential causality between mechanical cues (matrix stiffness) and cellular (energy) metabolism remains largely unexplored. Here, we culture human mesenchymal stem cells (hMSCs) on stiff (20 kPa) or soft (1 kPa) substrate and demonstrate that cytoskeletal reorganization and FA formation spreading on stiff substrates lead to a drop in intracellular ATP levels, correlates with the activation of AMP-activated protein kinase (AMPK). The resulting increase in ATP levels further facilitates cell spreading and reinforces cell tension of the steady state, and coincides with nuclear localization of YAP/TAZ and Runx2. While on soft substrates (1 kPa), lowered ATP levels limit these cellular mechanoresponses. Furthermore, genetic ablation of AMPK lowered cellular ATP levels on stiff substrate and strongly reduced responses to substrate stiffness. Together, these findings reveal a hitherto unidentified relationship between energy expenditure and the cellular mechanoresponse, and point to AMPK as a key mediator of stem cell fate in response to ECM mechanics.

## 1. Introduction

The physical properties of the extracellular matrix (ECM) have a profound impact on cell behavior and stem cell fate, and a direct relationship between matrix stiffness, cell spreading and lineage selection has been demonstrated [1, 2, 3, 4, 5, 6]. Forces generated within the actin cytoskeleton and transmitted through FAs, play a major role in the cellular response to biophysical cues [7, 8, 9, 10]. Much attention has been given to how cells respond to mechanical forces. However, less is known how these forces are integrated with other cellular processes including bioenergetics [11, 12, 13]. In order to understand how mechanical forces shape the cellular phenotype and regulate cell fate, it is necessary to consider the highly dynamic nature of the cellular response, which involves a hitherto overlooked large energy expenditure. Growing evidence indicates that the transduction of external mechanical forces is linked to metabolic signals such as neutral lipid synthesis [13], mitochondrial structural remodeling [14], cellular glucose uptake [11] and cell migration [15]. The cellular response is a well-tuned process that must balance energy supply with energy demand to allow proper actin polymerization, FA formation and cellular contractility buildup. But it remains unclear how cells respond to this energy expenditure, tune energy supply and demand, and maintain energy homeostasis for mechanotransduction.

AMPK, a well-characterized cellular energy sensor [16], plays a key role in the coordination of cell function by controlling intracellular ATP levels [17]. AMPK is Thr172-phosphorylated and thereby activated (yielding pAMPK) when the AMP/ATP ratio increases. This occurs for instance during starvation, hypoxia or cell detachment from the ECM [18, 19, 20]. Upon AMPK activation, energy homeostasis is restored by altering glucose, protein and lipid metabolism, paralleled by mitochondrial morphology changes [21]. Mechanical cues, chemical stimulations or metabolic stresses [22, 23, 24] can all lead to alterations in mitochondrial morphology, and AMPK is considered an important regulator of mitochondrial dynamics through phosphorylation of mitochondrial fission factor (Mff), thereby stimulating mitochondrial fission [24, 25]. A recent study in MCF10A (human breast epithelial) cells and MDCK II (canine kidney epithelial) cells, revealed that external forces from neighboring cells triggered AMPK activation via E-cadherin to increase cellular ATP levels and enhance actomyosin contractility, thus linking energy homeostasis with cell-cell adhesion mechanotransduction [11]. Actomyosin contractility also plays a major role in the cellular response to the physical properties of the ECM through FA formation and organization of the actin cytoskeleton, which are ATP-demanding processes [11, 12, 26].

In this study, we aim to elucidate the role of energy expenditure in the cellular response to substrate stiffness. When cells were seeded on stiff substrates, we observed an initial drop in ATP levels followed by higher ATP levels at 24 hrs, increased glucose uptake and actomyosin contractility, altered mitochondrial morphology, and sustained AMPK activation, resulting in nuclear localization of YAP/TAZ and Runx2, and osteogenic differentiation. Independent of substrate stiffness, cell fate appeared strongly correlated with the activation or inhibition of AMPK. Our findings establish a critical role for AMPK in connecting intracellular energy expenditure and cellular mechanotransduction, with downstream effects on stem cell fate.

## 2. Materials and Methods

### Preparation of polyacrylamide (PAAm) hydrogels

This method was adapted from a previously described procedure [4, 5]. 13-mm-diameter borosilicate glass coverslips (VWR) were freshly oxidized by oxygen plasma and then incubated in a solution composed of 0.3% w/v 3- (trimethoxysilyl)propyl methacrylate (Sigma Aldrich) and toluene (Fisher Scientific) overnight. Slides were washed with ethanol and dried with nitrogen. The stiffness of PAAm gels was regulated through the concentration of acrylamide (AA) and bis-acrylamide (BA). PAAm gel solutions were prepared with AA at final concentrations of 8, 20 and 30% w/v and BA at 0.02, 0.15 and 0.375% w/v. 5 μL of 10% w/v ammonium persulfate (Sigma Aldrich) and 1.5 μL TEMED (Sigma Aldrich) were added to AA/BA solutions to initiate PAAm polymerization. 4 μL of PAAm solution was immediately pipetted onto the pre-treated 13-mm-diamter coverslips and covered with an untreated 20-mm-diameter coverslip. The polymerization was completed in about 2 h, and then the samples were immersed in PBS buffer overnight. The top coverslips were carefully peeled off to obtain the gels adhering to the bottom coverslips. With the different concentrations of AA/BA described above, PAAm gels with different stiffness can be prepared. 8% w/v AA and 0.02% w/v BA, 20% w/v AA and 0.15% w/v BA, 30% w/v AA and 0.375% w/v BA showed stiffness of ~ 1 kPa, 20 kPa and 100 kPa, respectively.

### PAAm gels functionalization with collagen I

For cell adhesion, rat tail collagen I (BD biosciences) was crosslinked to the PAAm gels using N-sulfosuccinimidyl-6-(4′-azido-2′-nitrophenylamino)hexanoate (sulfo-SANPAH, from Life Technologies). 30 μL of 1mg/mL sulfo-SANPAH dissolved in milliQ H2O was added to PAAm gels which were irradiated under a 365nm UV lamp (ABM, USA) for 5 min. Then, gels were washed twice with PBS and the crosslinking procedure was repeated once more. After that, the samples were soaked in 50 μL/mL collagen solution for 2 h at room temperature. Finally, samples were washed twice with PBS before cell seeding.

### Cell culture

hMSCs, WT mouse embryonic fibroblasts (MEFs), AMPKα-null MEFs and NIH-3T3 mouse fibroblasts were cultured in low glucose DMEM (2 g/L, from Gibco), with 10% filtered fetal bovine serum (FBS, from Gibco), 1% penicillin-streptomycin (pen/strep, from Thermo) and 1% glutamine (Gibco). hMSCs were obtained from Lonza and cultured to passage 6 before seeding onto gels at a low density of 1250 per cm^2^ for studies involving cell spreading, mitochondrial dynamics, ATP measurements or specific proteins staining, 5000 per cm^2^ for glucose uptake, 2500 or 25000 per cm^2^ for osteogenic differentiation on stiff or soft gels, respectively, and 25000 per cm^2^ for adipogenic differentiation. For hMSCs cultured on gels, proliferation medium containing high glucose DMEM (4.5 g/L, from Gibco), 10% FBS, 1% glutamine and 1% pen/strep was used. For energy starvation, cells were cultured in DMEM without glucose, pyruvate or glutamine (Gibco) supplemented with 10 % FBS. For osteogenic or adipogenic differentiation, differentiation medium contained of proliferation medium and osteogenic/adipogenic chemical supplements (5×10^-7^ M dexamethasone, 5 mM β-glycerolphosphate, 0.1 mM ascorbic acid-2-phosphate, 250 μM 3-isobutyl-1-methylxanthine, 5 μg/mL insulin, and 5×10^-8^ M rosiglitazone maleate, all from Sigma). For tests of F-actin inhibition (2.5 μM Cytochalasin D, from ChemCruz), myosin inhibition (50 μM Blebbistatin, from Calbiochem), Rock inhibition (50 μM Y27632, from Sigma), AMPK inhibition (Compound C, from Sigma) or AMPK activation (A-769662, from Cayman Chemical), these chemicals were added to the proliferation medium or differentiation medium after 10 - 30 min of cell culture. WT MEF and AMPKα-null MEF were gifts from Benoit Viollet (Paris Descartes University), they were cultured and tested as described previously [27]. 5000 per cm^2^ of cells were seeded onto gels for all following experiments including ATP measurements, glucose uptake, morphology visualization and mechanotransduction markers staining.

### Cell staining

After incubation, cells were fixed with 4% paraformaldehyde (PFA) (Sigma) for 10 min followed by washing three times with PBS and permeabilized with 0.2% Triton X-100 (Sigma) for 10 min at room temperature. For F-actin and G-actin visualization, cells were stained with phalloidin-Atto 633 (1:1000, from Sigma), Alexa fluor488–DNaseI (1:500, D12371, from Invitrogen), 4’, 6-diamidino-2-phenylindole (DAPI) (1:1000, from Millipore) for 1 h. For vinculin, RhoA, β-integrin, myosin, pAMPK, YAP/TAZ and Runx2 staining, nonspecific binding sites were blocked in 10% BSA solution for 1 h, followed by incubation with primary antibody anti-vinculin (1:500, AB18058, from Abcam), anti-integrin β1 (1:500, AB30394, from Abcam), anti-RhoA (1:500, AB187027, from Abcam), anti-myosin IIa (1:500, 150M4764, from Sigma), anti-AMPK alpha 1 (phospho T183) + AMPK alpha 2 (phospho T172) (1:500, AB23875, from Abcam), anti-YAP/TAZ (1;500, D24E4, from Cell Signaling) and anti-Runx2 (1:500, AB76956, from Abcam) for 1 h. Subsequently, stained with DAPI, phalloidin and secondary antibody, Alexa-488 goat antimouse (1:1000, A11029, from Thermo Fisher Scientific) or anti-rabbit IgG (1:1000, A10040, from Thermo Fisher Scientific) for 1 h at room temperature.

### ATP assay

Intracellular ATP levels were measured using ApoSENSOR™ ATP Cell Viability Bioluminescence Assay Kit (Biovision), according to the manufacturer’s instructions. Briefly, 2500 cells were plated onto 13-mm-diameter gels (n≥6 per condition) and cultured to specific time points. Cells were lysed with 100 μL of Nuclear Releasing Buffer for 5 min at room temperature with gentle shaking. For the ATP levels at the starting time point (0 h), 2500 cells in suspension were collected and lysed directly. Then adding 10 μL ATP Monitoring Enzyme to the lysate and reading the sample within 2 min in a luminometer (Multimode Microplate Reader, from Berthod). The lysed cells were stained with DAPI and imaged with a Leica SP8 confocal microscope. Cell number on each gel was calculated from DAPI staining, it was used to normalize the ATP intensity to ensure an equal number of cells.

### Glucose uptake

Glucose uptake assays were performed by Glucose Uptake Cell-Based Assay Kit (Cayman Chemical) following the manufacturer’s instructions. This kit employs 2-NBDG, a fluorescently-tagged glucose derivative, as a probe for the detection of glucose taken up by cultured cells. Cells were plated onto 13-mm-diameter gels (n≥6 per condition) and cultured to 20 h on substrates with different stiffness or with different cell culture medium. Immediately following this, cells were treated in glucose-free medium for 2 h. At the end of treatment, 2-NBDG was added to a final concentration of 200 μg/mL in glucose-free medium for 30 min incubation. For fluorescence images, samples were visualized by Leica SP8 immediately. For uptake intensity test, the medium was carefully aspirated without disturbing the cell layer and the Cell-Based Assay Buffer was added to cells. After lysing cells, 100 μL of the supernatant was collected to a 96-well plate immediately. An additional 50 μL Cell-Based Assay Buffer was added, followed by readout at 485/535nm via microplate reader (Berthold). The number of cells was quantified by DAPI staining to normalize the glucose uptake.

### Proliferation assay

EdU labeling, which can incorporate into the DNA of cells during replication, was performed for proliferation studies. hMSCs were seeded on gels with different stiffness or treated with AMPK inhibitor or activator for 20 h, followed by treatment with EdU solution. When the cells were cultured to 48 h, followed by fixed and permeabilized with 4% PFA and 0.1% Triton X-100, respectively. Subsequently, samples were treated according to the manufacturer’s protocol of Click-iT EdU Alexa Fluor-488 HCS Assay (Thermo Fisher Scientific). Samples were imaged by Leica SP8 confocal microscope (Leica, Germany) with 10× objective to collect regions of interest.

### Mitochondrial imaging

For time-lapse imaging, hMSCs were cultured on gels with different stiffness and placed in a 24 well glass bottom plate (In Vitro Scientific), then mitochondria were stained with 100 nM MitoTracker Deep Red (Life Technologies). Up to 30 min incubation, a Leica SP8 microscope was used to track the mitochondrial dynamics. Individual images were acquired at a 2 min interval with a 20× objective at 37 °C and 7.5% CO_2_ atmosphere. After acquisition, images were analyzed using the Image 5D plugin of Fiji and exported as uncompressed AVI sequences. For mitochondrial visualization, 30 min prior to imaging at specific time points after seeding, Mito Tracker Deep Red was added to the culture medium. Cells were fixed with 4% PFA, permeabilized with 0.2% Triton X-100 and then counterstained with DAPI and phalloidin tetramethyl-rhodamine B isothiocyanate (TRITC) (Millipore). Fifteen-image Z-stacks were acquired by SP8 confocal with a 63× oil-immersion objective and merged with Fiji software (http://fiji.sc/).

### Mitochondrial membrane potential

The membrane potential was determined by TMRM Mitochondrial Membrane Potential Assay Kit (Abcam). Cells were cultured on the gels with different stiffness for 20 h, and then adding 100 nM TMRM to the medium 20 min before imaging. Live cells were immediately visualized without removing TMRM dye at 37 °C and 7.5% CO2 atmosphere and images were captured by the 549/575 nm laser of a Leica SP8 confocal microscope with a 40× water immersion objective at the photon counting mode.

### Western blotting

Cells were lysed with RIPA buffer (1:10, Cell Signaling) supplemented with protease and phosphatase inhibitors (Roche). Protein were collected and quantified using the BCA method. Equal amounts of proteins were loaded and separated on 12% Criterion Precast Gel (Bio-Rad), subsequently transferred onto PVDF membranes (Bio-Rad) and blocked with 5% non-fat dry milk (Bio-Rad) for 1 h at room temperature. AMPK and pAMPK were stained using primary antibody polyclonal AMPK (1:1000, 2532, from cell signaling) and anti-phosphorylated AMPK (Thr 172) (1:1000, 2535, Cell Signaling), β-actin (1:2000, from Abcam) was performed as loading controls and then incubated overnight on the roller bank at 4 °C. HRP-conjugated secondary antibodies, anti-rabbit or anti-mouse immunoglobulins (1:3000, Dako), were applied for 1 h at room temperature and detected using SuperSignal chemiluminescent substrates (Pierce).

### Differentiation assay

hMSCs were cultured for 7 or 10 days in mixed medium for osteogenic and adipogenic differentiation, respectively. The mixed medium containing the AMPK inhibitor or activator was changed twice a week. After incubation up to 7 or 10 days, all samples were fixed with 4% PFA and penetrated with 0.2% Triton-X 100 for 10 min, respectively. Osteogenic differentiation was analyzed by alkaline phosphatase (ALP) staining with Fast Blue assay (naphthol-AS-MSC phosphate and Fast Blue RR, Sigma) in Tris-HCl buffer (pH 8.9) and incubating at 37 °C for 1 h. Adipogenic differentiation was analyzed by Oil Red O staining with incubating cells with 1.8 mg/mL Oil Red O (Sigma) for 30-60 min at room temperature and then washing with 60% isopropanol (Sigma). Finally, nuclei visualization was performed by DAPI and images were collected by Zeiss inverted microscope (Photometrics, USA). The quantification of osteogenic differentiation was performed by manually counting the number of ALP positive cells in relation to the number of DAPI stained cells.

### Microscopy image analysis

Fluorescence images were acquired using a Leica SP8 confocal microscope with different objectives and overlaid in Fiji software with Image 5D plugin. All analysis of cells was based on single cells not in contact with other cells. For quantification of fluorescence intensity, images were taken with photon counting mode to make sure all settings were fixed. For quantification of G/F actin ratio, it was performed as our previous studies [28, 29]. The integrated fluorescence of F-actin and G-actin z-stack images was measured for individual cells, followed by subtraction of the background fluorescence. For quantification of YAP/TAZ and Runx2 subcellular localization, nuclear localization was defined as a ratio of the average intensity of target proteins in the nucleus compared to the average intensity of target proteins in the cytoplasm was larger than 1.

Mitochondrial morphology was quantified in cells stained with MitoTracker Deep Red using Image Pro Plus^®^ software (Media Cybernetics, Rockville, MD, USA) and an algorithm described in detail previously [30]. In brief, microscopy images were converted to 8-bit grayscale and background-corrected to obtain a COR image. The COR image was contrast-optimized using a linear contrast stretch (LCS) operation. Next, the LCS image was subsequently processed using a top-hat filter (THF) and median filter (MED). Then, the MED image was thresholded to obtain a binary (BIN) image, representing mitochondrial objects in white on a back background. Finally applying a Boolean AND operation on the COR and BIN image yielded a masked (MSK) image, in which mitochondrial area (*Am*; in pixels; a measure of mitochondrial size) and form factor (*F*; a combined measure of mitochondrial length and degree of branching) were quantified. Mitochondrial morphology measurements were carried out in 3 independent experiments in each of which >25 cells were analyzed.

### Statistical analysis

Statistical analysis was performed with Origin 2018 to assess the significance between data. One-way ANOVA tests was applied to compare data between multiple groups with a Tukey post-test and Student’s t-test was used for two variables. Significant differences are indicated by * P < 0.05 or **P < 0.01. NS indicates no significant difference (P > 0.05). All results are presented as mean ± standard error. All experiments were performed at least three independent experiments or biological replicates. The number of independent experiments (N) and the number of data points (n) in each experiment are indicated within corresponding figure legends.

### Data availability

The authors declare that the data is presented within the paper and its Supplementary Information or will be made available by the corresponding author upon reasonable request.

## 3. Results

### 3.1 Feedback between spreading and intracellular energy expenditure

We first determined the influence of substrate stiffness on indicators of energy expenditure such as intracellular ATP levels and glucose uptake rates. We cultured hMSCs on 1 kPa (soft) or 20 kPa (stiff) polyacrylamide (PAAm) gels functionalized with collagen (50 μg/mL), and determined intracellular ATP levels at different time points, from initial adhesion and spreading of cells to steady state (when no further morphological changes were observed) (**Figure 1a and Supplementary Figure 1a, b**). Unless noted, cells were always seeded at low density (1250 cells/cm^2^) to avoid contributions form cell-cell interactions. Consistent with previous findings [1, 5], cells grown on stiff substrates displayed larger cell spreading areas with reinforced F-actin stress fibers and accompanying lower levels of G-actin, and greater sizes and numbers of FAs (as determined by Vinculin staining) (**Supplementary Figure 1c-e**). During the same 3 h time window after seeding, intracellular ATP levels on the stiff substrate decreased by ~27% (**Figure 1a**), consistent with increased energy expenditure for spreading cells and concomitant formation of actin cytoskeleton and FAs. At steady state (no further changes in spreading area after 20 h), intracellular ATP levels on stiff substrates recovered and even exceeded those on soft ones (**Figure 1a**). Similarly, mouse embryonic fibroblasts (MEF) and NIH 3T3 cells also displayed increased intracellular ATP levels when cultured for 20 h on substrates of increasing stiffness (**Figure 1b)**. As glucose is the primary source of cellular energy production in hMSCs [31], we examined if different substrate mechanics result in changes in glucose uptake. We observed a 31% increase in glucose uptake in hMSCs cultured on 20 kPa compared to 1 kPa PAAm substrates as demonstrated by the higher intracellular levels of 2-NBDG, a fluorescent glucose analog used for monitoring glucose uptake into living cells (**Figure 1c)**. A correlation between energy demand and spreading was also evident in glucose starvation experiments. Reducing glucose concentration in culture medium from 4.5 to 0 g/L for cells on stiff substrates significantly impacted on intracellular ATP levels (~34% lower; **Figure 1d**) and starvation halted cell proliferation (**Supplementary Figure 2**). To investigate the role of energy in more detail, we monitored hMSC spreading on stiff substrates in the presence or absence of glucose **(Figure 1e and Supplementary Figure 1b, 3)**. Removal of glucose from the culture medium resulted in significantly reduced actin polymerization and FA formation, reflected by a 3.8-fold increase in G/F-actin levels (**Figure 1f**) after 20 h culture. We further studied the temporal characteristics of intracellular ATP changes upon disruption and re-organization of the actin cytoskeleton. To do this, cells were treated for 1 h with Cytochalasin D (CytoD) [8], which disrupts the actin cytoskeleton, and resulted in cells displaying a small rounded morphology (**Figure 1g**). After CytoD removal from the medium, cells were allowed to re-spread on the substrate, during which intracellular ATP levels dropped by ~22% (**Figure 1g**), This strongly suggests that the actin cytoskeleton organization consumes a significant fraction of intracellular ATP. Both actin polymerization and FA formation lead to increased actomyosin activity. To investigate if this activity also impacted on intracellular ATP level, cells were cultured on stiff substrates and treated with two inhibitors of actomyosin contractility: Rho-associated protein kinase (ROCK) inhibitor (Y-27632) and myosin inhibitor blebbistatin (Bleb.) [32]. Instead of the observed drop in ATP levels earlier, treatment with inhibitors resulted in stable cellular ATP levels (normalized to the obtained values at 0 h) consistent with a less developed actin cytoskeleton and reduced cell contractility (**Figure 1h and Supplementary Figure 4**).

**Figure 1.**
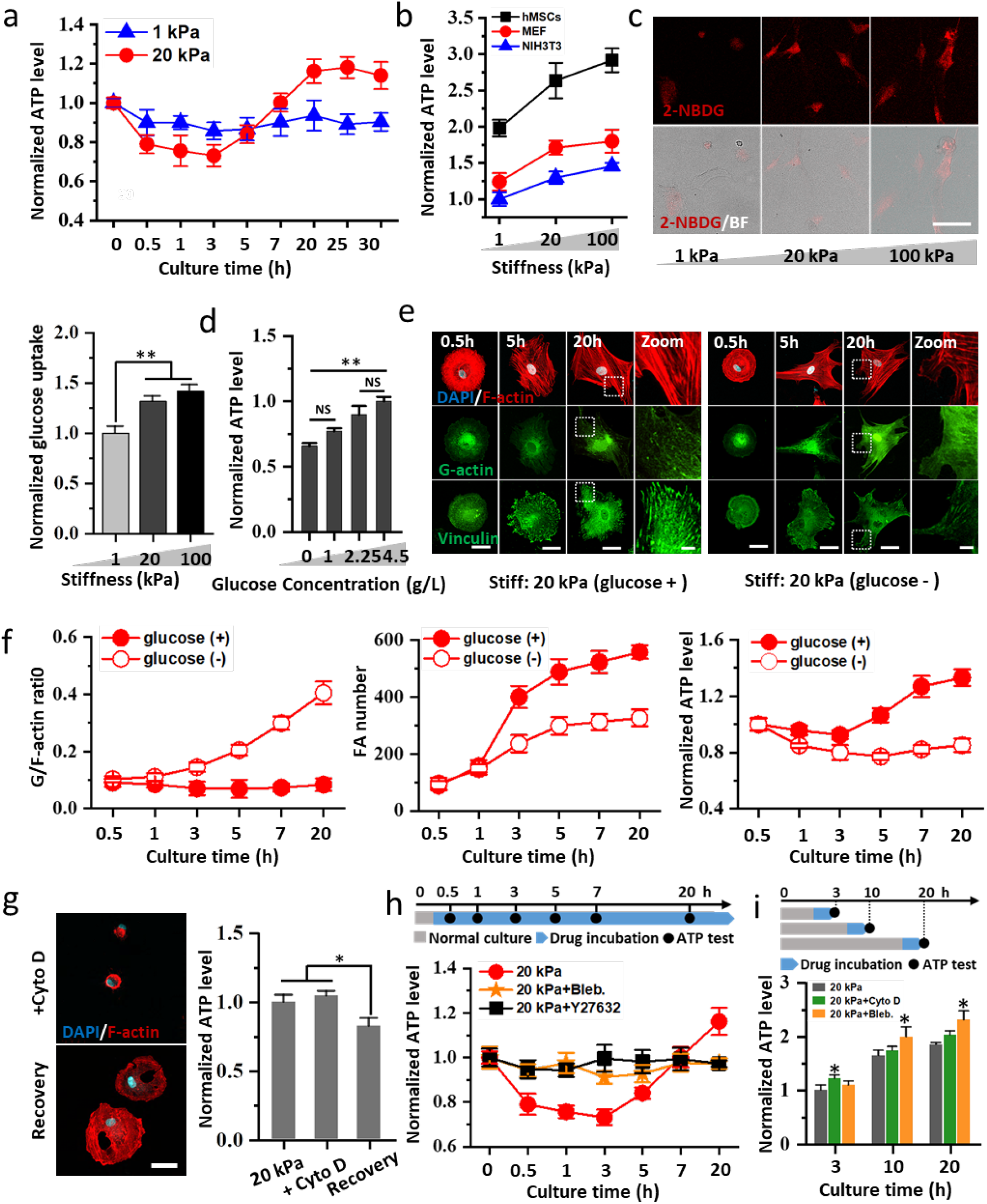
Intracellular ATP levels respond during cell spreading and adhesion. (a) Normalized ATP levels of hMSCs cultured on soft (1 kPa) and stiff (20 kPa) PAAm gels over time (n=6 biologically independent samples, N=3). Intracellular ATP level at time 0 was measured from cells in suspension (before seeding on PAAm hydrogel) (b) ATP levels in different types of cells (hMSCs, MEF, NIH3T3) at 20 h after seeding on PAAm gels (1, 20 and 100 kPa) (n=6 biologically independent samples, N=3). (c) Glucose uptake on different substrates visualized by 2-NBDG (red). Graph: quantification of 2-NBDG intensity (n=5 biologically independent samples, N=3). Scale bars, 100 μm. (d) Intracellular ATP levels in response to different glucose concentration in the medium (n=6 biologically independent samples, N=3). (e) Confocal images of hMSCs cultured in the presence or absence of glucose on stiff PAAm gels showing G-actin (stained via labeled DNaseI: green) and FAs (vinculin staining: green) at different time points. F-actin was stained with phalloidin (red) and nuclei were counterstained with DAPI (blue). Scale bars, 50 μm (left three) and 10 μm (right). (f) Quantification of G/F actin ratio (n>75 cells from three independent experiments), FA numbers (n>75 cells from three independent experiments) and normalized ATP levels (n=6 biologically independent samples, N=3) in hMSCs cultured under glucose starvation on stiff PAAm gel. (g) Actin disassembly with 2.5 μM Cytochalasin D (Cyto D) and actin recovery after removal of Cyto D. Confocal images show the changes in cellular morphologies. Graph shows changes in intracellular ATP intensity (n=6 biologically independent samples, N=3). Scale bars, 50 μm. (h) Normalized ATP intensity in hMSCs cultured on stiff PAAm gel treated with 50 μM myosin inhibitor Blebbistatin (Bleb.) or 50 μM Rock inhibitor Y27632 (n=6 biologically independent samples, N=3). (i) Quantification of intracellular ATP intensity at 3, 10 and 20 h after 1 h of treatment with Cyto D and Bleb. (n>6 biologically independent samples, N=3). Images of (c, e, g) are representative of at least three independent experiments. Data are shown as mean ± s.d., * and ** indicate P values of < 0.05 and <0.01, respectively.

As cell spreading is a dynamic process, we wondered if disruption of actin cyctoskeleton formation and cell contractility at different time points would lead to different responses. We hypothesized that during the early spreading stage, actin polymerization would consume most ATP, whereas at steady state (i.e. in fully spread cells), maintaining cell tension would require higher energy expenditure than actin polymerization. To validate our hypothesis, we studied the relative contributions by inhibiting actin polymerization and cytoskeletal tension using CytoD and Bleb., respectively, at different time points (**Figure 1i**). For ease of comparison, ATP intensities were normalized to the non-treated control group at 3 h after seeding. At early time points (3 h), inhibition by CytoD restored intracellular ATP levels (up 1.2 fold); whereas Bleb. treatment had no significant effect. In contrast, at later time points (10 h and 20 h), treatment with CytoD did not lead to increases in ATP levels, but the addition of Bleb. did, showing a 12 % increase. These results indicate that during early stages of cell spreading, ATP expenditure due actin polymerization is indeed more important that cell tension, whereas at steady state the opposite is the case.

### 3.2 AMPK activation correlates with altered mitochondrial morphologies

In stem cells, mitochondria are prime generators of ATP [14, 24], these organelles are motile, and continuously fuse and divide, a process influenced by mechanical cues [14, 22]. In addition to its role in ATP production, mitochondrial morphology has been coupled to the coordination of self-renewal vs. differentiation of stem cells [33]. Primed by these findings we next determined whether substrate stiffness affected mitochondrial morphology. Visual inspection suggested that, on soft substrates, mitochondria display a filamentous structure at both 5 h and 20 h after seeding. In contrast, mitochondrial morphology appeared more fragmented on stiff substrates (**Figure 2a and Supplementary Figure 5**). Quantitative analysis [30] revealed that the mitochondrial area (*Am*, a measure of mitochondrial size) and form factor (*F*, a combined measure of mitochondrial length and degree of branching), were significantly reduced on stiff relative to soft substrates (**Figure 2b, c**). Mitochondria in cells on stiff substrates displayed a stronger staining with the fluorescent cation tetramethylrhodamine methyl ester (TMRM), (**Supplementary Fig. 6**), suggesting that the mitochondrial membrane potential of these mitochondria is more negative. Finally, time-lapse imaging of mitochondria suggested that the mitochondrial fission process was faster on stiff substrates compared to soft substrates. (**Supplementary Figure 7** and **Supplementary Movie**).

**Figure 2.**
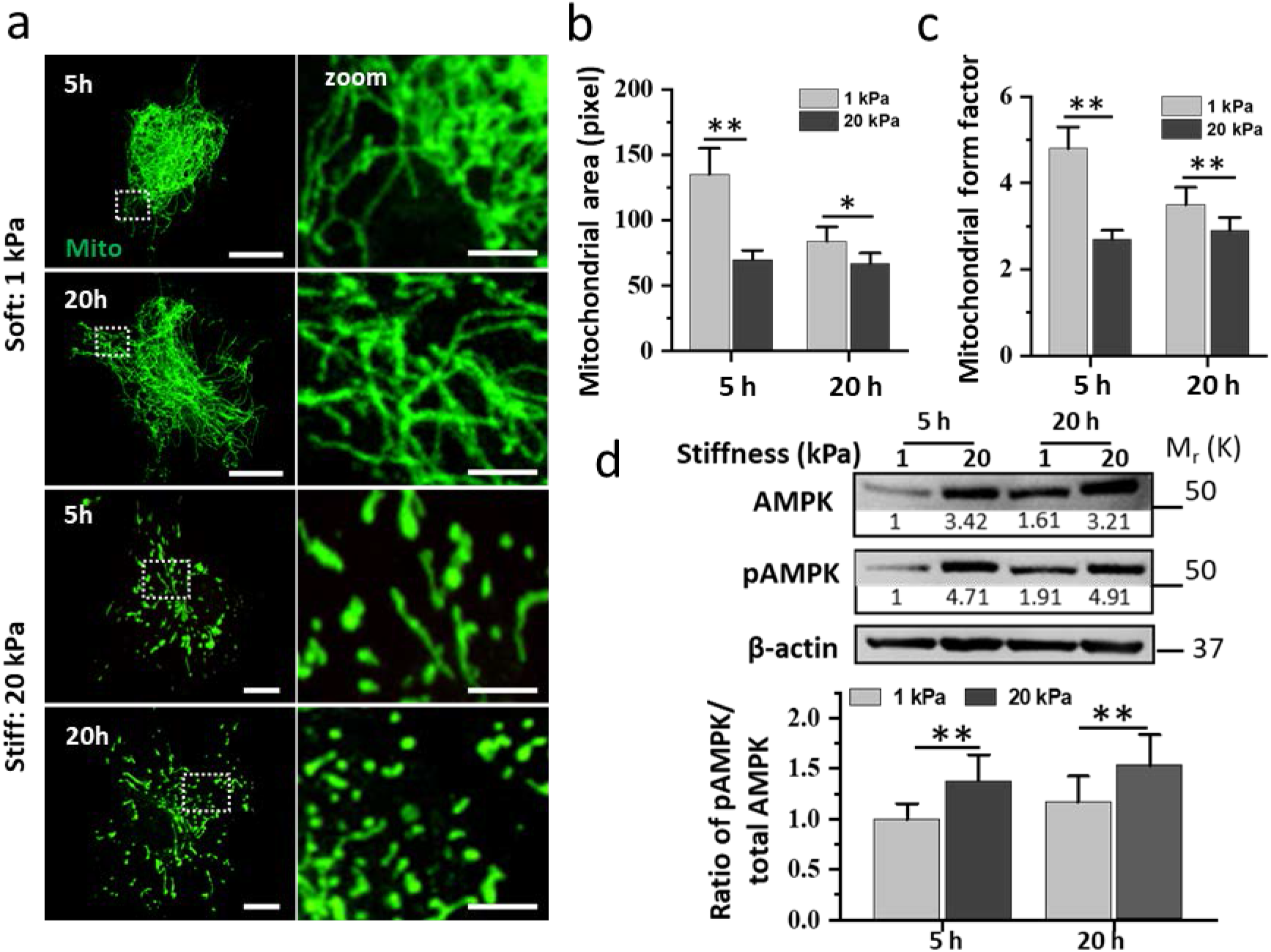
Changes in mitochondrial morphologies and pAMKP/AMPK ratios correlate with ECM stiffness. (a) Mitochondrial morphologies in hMSCs on soft or stiff PAAm gels at 5 or 20 hours after seeding. Mitochondria were stained with Mito tracker (green). Arrows refer to the zoomed region. Scale bars, 25 μm (left) and 10 μm (right). (b, c) Quantification of (b) mitochondrial area and (c) form factor of the cells shown in (b) (n>25 representative cells from three independent experiments). (d) Western blots of AMPK and Thr172-phosphorylated AMPK (pAMPK) after 5 or 20 h incubation on different PAAm gels. Graph: Quantification of pAMPK/AMPK ratios. AMPK and pAMPK were first normalized to actin (numbers shown on the blot) before determining pAMPK/AMPK ratios. Images in (a) are representative of at least three independent experiments. Data are shown as mean ± s.d., * and ** indicate P values of < 0.05 and <0.01, respectively.

It has been shown that AMPK mediates mitochondrial fragmentation in response to energy stress [24]. In this context, we hypothesized that the observed drop in ATP levels during cell spreading would result in a similar energy stress, leading to AMPK activation and induction of a fragmented mitochondrial phenotype. Western blot analysis revealed increased pAMPK/AMPK ratios in hMSCs cultured for 5 and 20 h on stiff vs. soft substrates (**Figure 2d**). To gain insight into the chain of events during AMPK phosphorylation we performed western blot and immunostaining analysis at different time points (**Supplementary Figure 8**). On stiff substrates, virtually no pAMPK was observed during initial cellular adhesion period (0.5 h) (**Supplementary Figure 8**). Cells cultured on these substrates for longer than 1 h displayed elevated pAMPK levels as well as nuclear localization. Together, these experiments establish that AMPK phosphorylation is preceded by a drop in ATP levels (**Figure 1a and Supplementary Figure 8**). Furthermore, a fragmented mitochondrial phenotype was observed 1 h after seeding (**Supplementary Figure 9**), in agreement with the time point at which increased pAMPK levels were observed.

### 3.3 AMPK plays a central role in mechanotransduction

Previous studies [11] have linked energy metabolism to E-Cadherin mechanotransduction as liver kinase B1 (LKB1) is recruited to the cadherin adhesion complex and activates AMPK. However, it is not known if AMPK plays a similar role in the mechanotransduction response from cell-matrix interactions. To further explore the role of AMPK in mechanotransduction, we cultured *AMPKα1^-/-^ AMPKα2^-/-^* (AMPK*α*-null) mouse embryonic fibroblasts (MEFs) on soft and stiff substrates. These cells displayed a 40-50% lower ATP content that was not affected by stiffness **(Figure 3a)**. In **Figures 3b, c**, a lower glucose uptake was shown in AMPK*α*-null MEFs on stiff substrates, compared to WT MEFs, as determined by 2-NBDG intensity. Interestingly, cell morphology of WT and AMPK*α*-null cells on soft substrates appeared very similar **(Figure 3d)**. In contrast, on stiff substrates AMPK*α*-null cells displayed limited spreading, and a less well-developed actin cytoskeleton (as evidenced by the ~2.3-fold higher G/F-actin ratio). **Figure 3e** demonstrates reduced actomyosin contractility in AMPK*α*-null cells on stiff substrates and a lower number of FAs. YAP/TAZ is a mechanical regulator that typically shows cytoplasmic or nuclear localization regulated by external signals [32, 34]. Stiff substrates and actomyosin tension are associated with high levels of nuclear YAP/TAZ [3, 35], and in WT cells we observed nuclear localization in 79±10% cells. Conversely, inhibition of actin polymerization and stress fiber formation by silencing AMPK almost halved YAP/TAZ nuclear localization **(Figure 3f)**.

**Figure 3.**
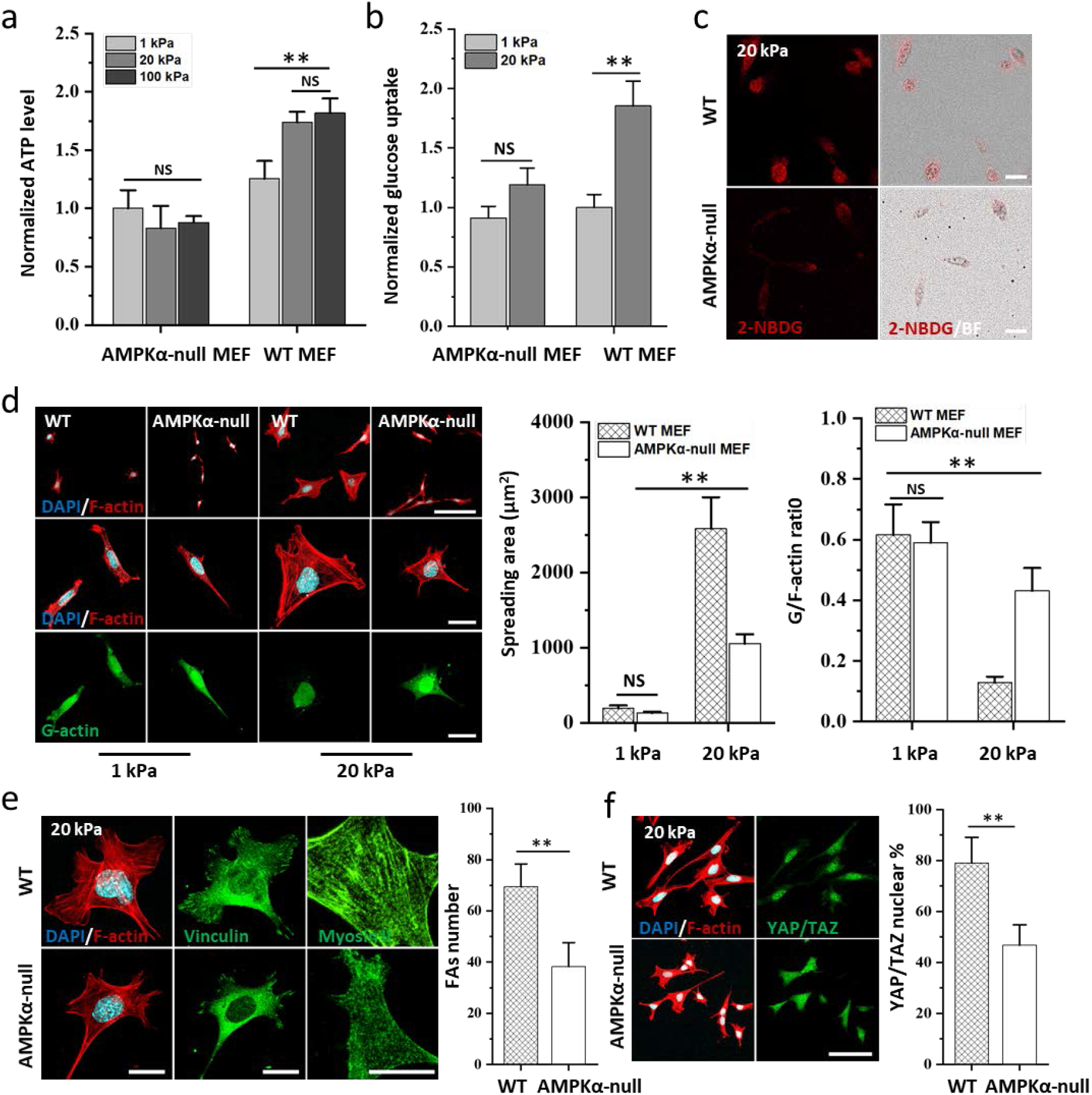
AMPKα-null cells show strongly reduced mechanoresponse on stiff substrates. (a) Normalized ATP levels in WT-MEF and AMPKα-null MEF cultured on 1, 20 and 100 kPa PAAm gels at 20 h after seeding (n=6 biologically independent samples, N=3). (b) Quantification of glucose taken up on different substrate (n=5 biologically independent samples, N=3). The difference in glucose uptake in WT MEF and AMPK-null MEF was compared between stiff and soft substrate. (c) Confocal images showing 2-NBDG (red) uptake on stiff substrate. Scale bars, 100 μm. (d) WT-MEF and AMPKα-null MEF cultured on stiff substrate demonstrating F-actin (red) and G-actin (green) after 20 h culture. Top: scale bar 100 μm; middle: scale bar 25 μm; bottom: scale bar 25 μm. Graph: quantification of spreading area and G/F-actin ratio (n>75 cells from three independent experiments). (e) Vinculin (middle: green) and Myosin II (right: green) expression in cells cultured on stiff substrate. Scale bar 25, μm. Graph: quantification of FA numbers (n>75 cells from three independent experiments) in cells cultured on stiff substrate. (f) YAP/TAZ localization in WT and AMPKα-null cells on stiff substrates. YAP/TAZ (green), F-actin and nuclei were counterstained with phalloidin (red) and DAPI (blue). Scale bars, 100 μm. Graph: Quantification of YAP/TAZ nuclear localization (n>120 cells from three independent experiments) in single cells on stiff substrate. Images of (c, d, e, f) are representative of at least two independent experiments with similar results. Data are shown as mean ± s.d., ** indicate P values of <0.01. NS indicates no significant difference with P > 0.05.

### 3.4 Manipulating intracellular AMPK activation alters cellular mechanoresponses

To further dissect the role of AMPK, we modulated its phosphorylation status using a cell-permeable AMPK inhibitor (Compound C) or AMPK activator (A-769662) at different concentrations after 30 min of cell spreading on stiff or soft substrates, respectively, and then evaluated changes in cellular response after culturing cells for 20h. It was found that increasing concentrations of AMPK inhibitor resulted in ~60% lower ATP levels on stiff substrates, while addition of AMPK activator on soft substrates led to 20% higher ATP levels (**Figure 4a**). The inhibition or activation of AMPK also impacted on glucose uptake, inducing a small (~13%) decrease after inhibition of AMPK on stiff substrates, and a 1.2~fold increase after activating AMPK on soft substrates (**Figure 4b**). Inhibition of AMPK on a stiff substrate induced a shift in mitochondrial morphology from a fragmented into a more filamentous phenotype. However, activation of AMPK on a soft substrate yielded a fragmented mitochondrial phenotype, previously associated with the cells cultured on stiff substrates (**Figure 4c and Supplementary Figure 10**). These experiments support a mechanism in which alterations in mitochondrial morphology are evoked by changes in AMPK activity. We next investigated whether inhibition or activation of AMPK was sufficient in regulating FA formation, cytoskeletal organization and YAP/TAZ localization. Inhibition of AMPK clearly influenced FAs, which appeared smaller and confined to the edge of spreading cells (**Figure 4d**). Furthermore, cytoskeletal organization was less developed, with fewer cross-cell stress fibers with increasing concentrations of AMPK inhibitor (**Figure 4d**). YAP/TAZ was increasingly localized in the cytoplasm after inhibition of AMPK (nuclear localization declined five-fold at 20 μM inhibitor concentration), while nuclear YAP/TAZ localization in cells on soft substrates increased upon AMPK activation (**Figure 4d**). Furthermore, we modulated energy substrate supply by varying the extracellular glucose concentration. YAP/TAZ nuclear localization only became pronounced at glucose concentrations >2.5 g/L and during glucose starvation YAP/TAZ nuclear localization was as low as 21±9% (**Supplementary Figure 11**). Nuclear YAP/TAZ localization induced by ECM stiffness requires stress fibers and cytoskeletal tension which can stretch nuclear pores and promote YAP/TAZ translocation. There is a balance of forces between cell adhesion on the outside and myosin II-based contractility on the inside of the cell, and myosin II thus plays an important role in controlling the mechanoresponse [36]. Inhibition or activation of AMPK clearly alters myosin levels (**Figure 4e**) and the changes mirror trends in YAP/TAZ localization.

**Figure 4.**
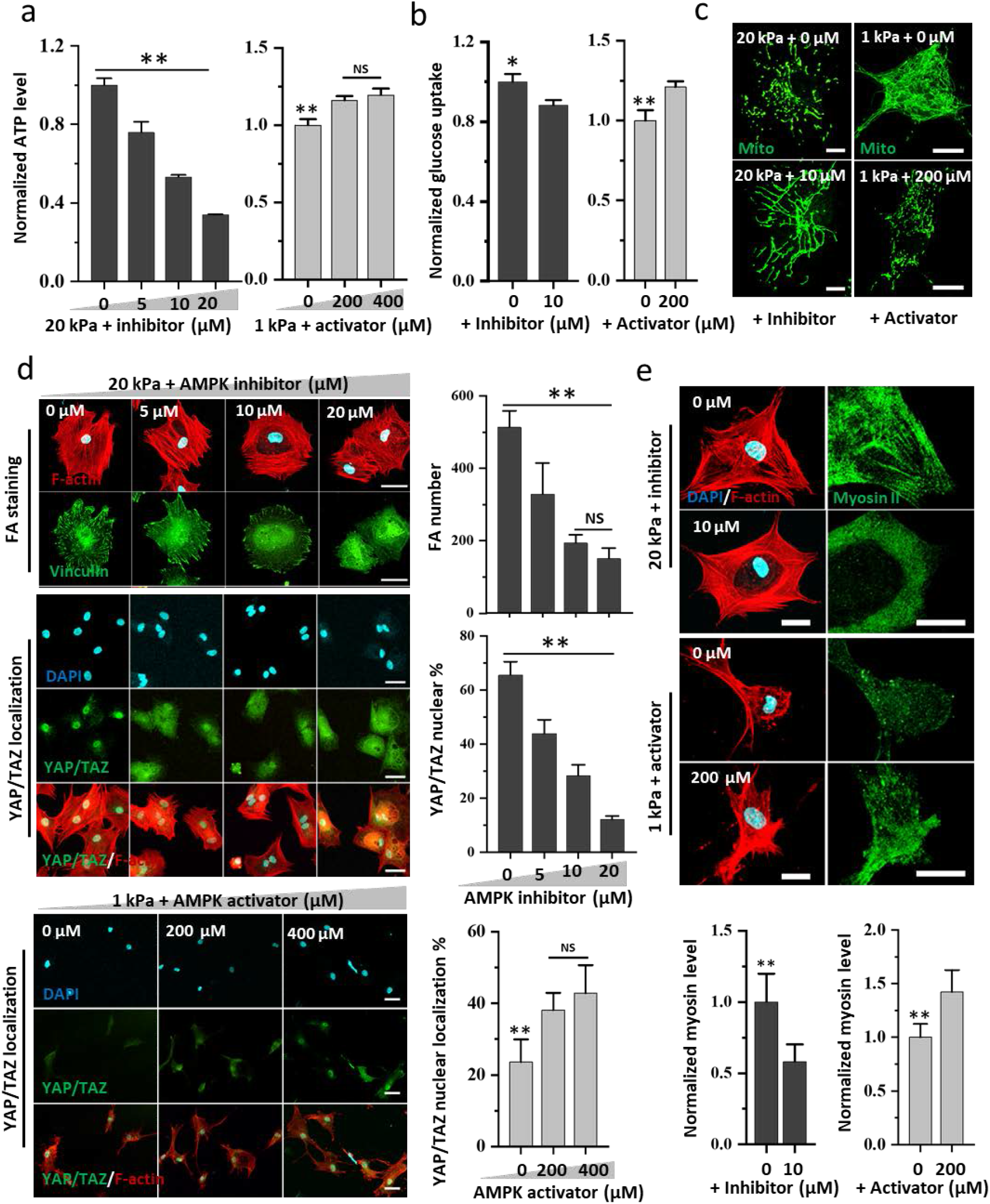
AMPK inhibition/activation alters ATP levels and cellular mechanoresponses. (a) ATP levels and (b) glucose uptake in hMSCs treated with variable concentrations of AMPK inhibitor (Compound C) on stiff substrate or treated with AMPK activator (A-769662) on soft substrates after 20 h culture (n=5 biologically independent samples, N=3). (c) Mitochondrial morphologies on stiff (left) or soft (right) substrates with (bottom) or without (top) AMPK inhibitor/activator treatment. Scale bars, 20 μm (left) and 10 μm (right). (d) The influence of AMPK inhibition/activation on cellular mechanoresponses. Confocal images of FAs (top) and YAP/TAZ localization (middle) in cells on stiff substrates treated with AMPK inhibitor, and YAP/TAZ nuclear localization in the presence of AMPK activator (bottom). Quantification of number of FAs (n>25 representative cells, N=3) and YAP/TAZ nuclear localization (n>120 cells from three independent experiments) in single cells. Scale bars, 50 μm. (e) The influence of AMPK inhibition/activation on cell tension. Expression of Myosin II in cells seeded on stiff (top two) or soft (bottom two) substrate treated with or without AMPK inhibitor/activator. Graph: quantification of Myosin II intensity of the cells (n>20 cells from two independent experiments). Scale bars, 25 μm. Images of (c, d, e) are representative of at least two independent experiments with similar results. Data are shown as mean ± s.d., ** indicate P values of <0.01. NS indicates no significant difference with P > 0.05.

### 3.5 AMPK activity influences stem cell differentiation into osteoblasts

Finally, we determined how manipulation of AMPK activity impacted on cell fate. Runx2 protein is a prominent transcription factor which can be detected in pre-osteoblasts and induces the differentiation of MSCs into osteoblasts when translocated into the nucleus of stem cells [37,38]. Compound C (10 μM) was added to the medium for cells cultured on stiff substrates and A-769662 (200 μM) was added to the medium for cells on soft substrates. Relative to standard culture conditions, a higher Runx2 nuclear localization was detected under conditions that promote higher AMPK activation on stiff substrates (*i.e*. with no inhibitor present) (**Figure 5a**) or additional AMPK activator on soft substrates (**Figure 5b**) after 1-day culture.

**Figure 5.**
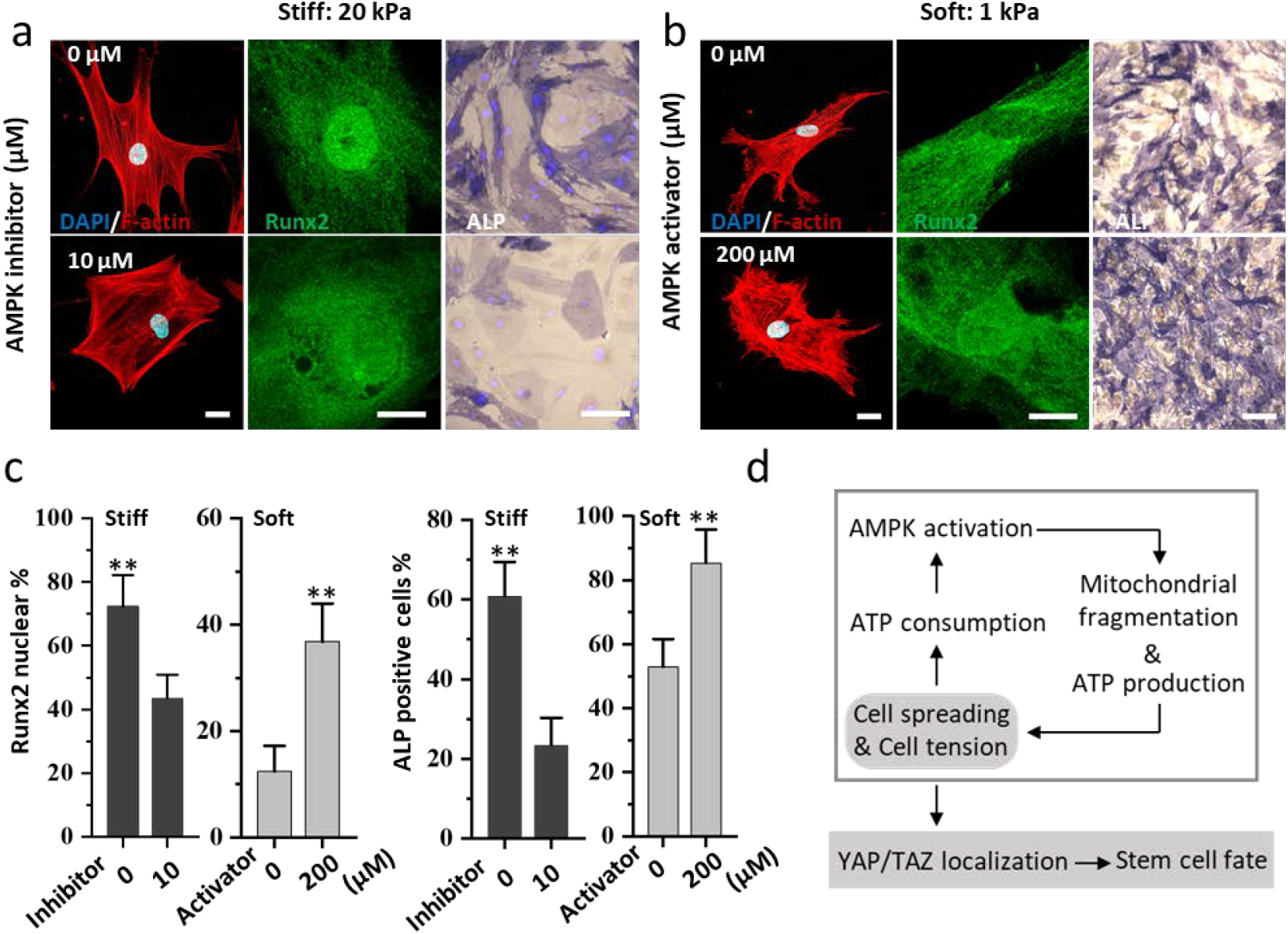
AMPK inhibition or activation affects stem cell differentiation. (a, b) Runx2 and alkaline phosphatase (ALP) staining showing osteogenic differentiation of hMSCs. Runx2 (green) localization and ALP (bright field) expression of the cells treated with (a) AMPK inhibitor on stiff substrates or (b) AMPK activator on soft substrates (bottom). Scale bars 20 μm (fluorescence images), 250 μm (bright field images). (c) Quantification of Runx2 nuclear localization after 1 day (n>30 cells from two independent experiments) and ALP positive cells after 7 days (n>6 representative images from two independent experiments) at different conditions. (d) Schematic overview showing the link between mechanotransduction and energy homeostasis. Images of (a, b) are representative of at least two independent experiments with similar results. Data are shown as mean ± s.d., ** indicate P values of <0.01. NS indicates no significant difference with P > 0.05.

Further, hMSCs were seeded on different substrates and cultured in mixed adipogenic/osteogenic differentiation medium. After 7-10 days culture time, ALP staining was used to indicate osteogenic differentiation and Oil Red O staining was performed for adipogenic differentiation. ALP staining showed AMPK inhibition in cells on stiff substrates significantly decreased ALP intensity, indicating lower osteogenic differentiation, whereas on soft substrates ALP staining was significantly enhanced in the presence of AMPK activator (**Figure 5c**; here higher cell densities were necessary, 25000 cells/cm^2^ instead of 2500 cell/cm^2^ on soft substrates, as cell numbers otherwise were too low to allow differentiation after extended culture times). The observation that AMPK modulation can override the impact of biophysical cues in the case of osteogenic differentiation correlates with the notion that such differentiation is typically seen on stiff substrates, where cells develop strong stress fibers and high intracellular tension. AMPK modulation affects these energy-demanding processes. Interestingly, we did not observe statistically relevant differences when studying adipogenic differentiation using Oil Red O staining when we either inhibited AMPK on stiff substrates, or activated AMKP on soft substrates (**Supplementary Figure 12**). Adipogenic differentiation is typically favored on soft substrates where cells have less pronounced stress fibers and associated energy demands, and this might explain why the modulation of AMPK does not lead to significant differences.

## 4. Discussion

Adhering and spreading cells form focal adhesions and a network of dynamic actomyosin stress fibers that maintains a certain tensional homeostasis adapted to the mechanical properties of the ECM [1, 5, 39]. These processes require energy, which raises the question of how cells ensure the availability of this energy and whether high energy expenditure limits cellular responses. An elevated ATP: ADP ratio due to F-actin formation [40], cytoskeleton remodelling [41], and adhesion mediated contractility [42] has been reported to increase glucose uptake during cell migration [43], the relationship between the cell dynamics and increased energy utilization. In this study we observed a clear link between mechanotransduction and the energy expenditure needed for the formation of well-developed FAs, cytoskeletal reorganization into well-defined stress fibers in spread cells, as well as the contractility of the cytoskeleton. Intracellular ATP levels are strongly affected by these energetically costly processes. In order to look into details about the extent of ATP consumption used for actin polymerization and contractility, we selectively inhibited actin polymerization by CytoD and actomyosin by Bleb. and revealed that inhibition of actin, but not myosin, restored intracellular ATP levels at early time points. However, at steady state (after cell area was at equilibrium, or better: steady state), inhibition of myosin, but not actin, restored intracellular ATP levels. Those experiments suggested that cells consume ATP to organize cytoskeleton at early times during spreading, to maintain tension at later time points (at steady state) and approximately 20% of cellular ATP is consumed by these processes.

A further link between energy metabolism and development of the actin cytoskeleton is highlighted by glucose starvation experiments. In the latter, glucose depletion significantly reduces actin polymerization and FAs formation, confirming that cellular mechano-responses consume a significant fraction of intracellular ATP. We demonstrate (**Figure 1**) that cell spreading on stiff substrates is associated with an initial drop in ATP levels. This drop activates AMPK, a kinase that is triggered by changing AMP/ATP ratios that arise from, for example, starvation, hypoxia or cell detachment from the matrix [18–20]. Recent studies also provided other AMPK activation pathways in various cell types, including Ca^2+^ influx [44, 45] or cell-cell forces [11]. Here we show that AMPK is activated coinciding with the ATP drop and mitochondrial fragmentation (**Figure 2**). As reported previously, mitochondrial trafficking to the leading edge of the cell could support the formation of protrusions and focal adhesions, necessary for cell migration and force generation [15, 46, 47]. Here, we showed that mitochondrial morphology was highly responsive to AMPK activation which is in full agreement with previous studies [24]. This strongly suggests that AMPK activation or inhibition directly affects mitochondrial morphology, indicating that changes in mitochondrial morphology were coupled to energy metabolism. Given the fact that AMPK plays a central role in the coordination of cell metabolism and function [17], its activation likely impacts on nuclear localization of YAP/TAZ. In this context, previous reports demonstrated that AMPK lowers YAP activity via phosphorylation [20], evoking the cytoplasmic localization of YAP/TAZ. Conversely, we find that YAP/TAZ is predominantly localized in the nucleus (**Figure 3f**), as typically observed in cells spreading on stiff substrates [3, 32]. It should be noted that in our study, cells were seeded at very low density to avoid cell-cell interactions, whereas previous work has shown that cell-cell contacts play a role in AMPK activation [11]. The main finding of our study is that maintaining a high cellular contractility is regulated (indirectly) by AMPK as high ATP levels are necessary to support actin polymerization, focal adhesions formation and actomyosin contractility. These processes set up a feedback loop that couples into YAP/TAZ localization. We present three different lines of evidence for placing YAP/TAZ translocation mechanistically downstream of AMPK activation: i) YAP/TAZ increasingly localized in the cytoplasm with declining ATP levels after inhibition of AMPK by Compound C treatment (**Figure 4d**). ii) AMPK activation was altered by changing the glucose concentration in the culture medium, and we found that YAP/TAZ nuclear localization only became pronounced at glucose concentrations >2.5 g/L, whereas glucose starvation resulted in YAP/TAZ nuclear localization as low as 21% (**Figure 4d**). iii) We found that ATP levels and glucose uptake were greatly reduced in a AMPKα-null mouse embryonic fibroblast (MEF) cell line, and YAP/TAZ localization in the nucleus on stiff substrates was also significantly reduced (by 32%). All these results firmly confirmed that AMPK activation can induce YAP/TAZ nuclear localization. We further observed that osteogenic differentiation appears correlated with activation or inhibition of AMPK, and not with the substrate stiffness.

In summary, our findings indicate that AMPK-mediated energy regulation plays an important role in the cellular mechanoresponse, as shown schematically in **Figure 5d**. The mechanoresponse on stiff substrates is initiated by cell spreading and the concomitant consumption of ATP to establish FAs and remodel the actomyosin network. Soon after initiation of cell spreading (i.e. in the first 0.5 h) ATP expenditure exceeds production, and the drop in intracellular ATP levels triggers activation of AMPK. This is consistent with recent evidence demonstrating that cells adapted their metabolic activity to variable mechanical cues through stress fiber formation and F-actin bundling [48]. After activation of AMPK, we observe a restoration of ATP levels, and ultimately (after 24h) even higher levels than before spreading. At the same time, we observe significant changes in mitochondrial morphologies pointing at increased ATP production. We found that upregulation of ATP production provides the required energy for further spreading and increased cell tension as the actin cytoskeleton organizes and FAs form, leading to a reinforcing cycle. Those experiments suggest that cells consume ATP to polymerize actin at early time points, and to increase cell tension at later time points. With the increased cell tension, cells that spread on stiff substrates then showed the expected YAP/TAZ nuclear localization and, ultimately, osteogenic differentiation.

## 5. Conclusions

Taken together, this work establishes temporal changes in intracellular ATP levels during cell spreading as a new mechanistic link between mechanical forces and molecular responses. In this sense, energy expenditure couples mechanical cues with cellular metabolism via AMPK activation in response to lowered ATP levels. These findings provide an impetus for further studies into the mechanisms that guide energy metabolic remodeling in cells with increased energy demands during their response to environmental cues. Of particular interest in this context is how the remodeled energy metabolism in tumor cells shapes their mechanoresponse to ECM alterations.

## Acknowledgments

We thank José M. A. Hendriks for assistance with cell cultures, and Dr. Liesbeth Pierson for assistance with confocal microscopy. We thank Dr. Benoit Viollet (Paris Descartes University) for providing WT-MEF and AMPKα-null MEF cell lines. J.X., M.B. and X.H. would like acknowledge the department of General Instruments of the Radboud University for providing confocal and light microscopy services. We acknowledge financial support from the China Scholarship Council (J.X. and X.H.) and the Radboud Nanomedicine Alliance (M.B.).

## References

[1] D. E. Discher, P. Janmey, Y. L. Wang, Science 2005, 310, 1139.

[2] A. J. Engler, S. Shamik, S. H Lee, D. E. Discher, Cell 2006, 126, 677.

[3] A. Elosegui-Artola, I. Andreu, B. Aem, A. Lezamiz, M. Uroz, A. J. Kosmalska, R. Oria, J. Z. Kechagia, P. Rico-Lastres, R. A. Le, Cell 2017, 171, 1397.

[4] B. Trappmann, J. E. Gautrot, J. T. Connelly, D. G. Strange, Y. Li, M. L. Oyen, M. A. C. Stuart, H. Boehm, B. Li, V. Vogel, Nat. Mater. 2012, 11, 642.

[5] T. Yeung, P. C. Georges, L. A. Flanagan, B. Marg, M. Ortiz, M. Funaki, N. Zahir, W. Ming, V. Weaver, P. A. Janmey, Cell Motil. Cytoskel. 2005, 60, 24.

[6] V. Vogel, M. Sheetz, Nat. Rev. Mol. Cell Biol. 2006, 7, 265.

[7] D. Choquet, D. P. Felsenfeld, M. P. Sheetz, Cell 1997, 88, 39.

[8] J. Adams, Cell. Mol. Life Sci. 2001, 58, 371.

[9] C. S. Chen, J. Tan, J. Tien, Annu. Rev. Biomed. Eng. 2004, 6, 275.

[10] J. D. Mih, A. Marinkovic, F. Liu, A. S. Sharif, D. J. Tschumperlin, J. Cell Sci. 2012, 125, 5974.

[11] J. L. Bays, H. K. Campbell, C. Heidema, M. Sebbagh, K. A. DeMali, Nat. Cell Biol. 2017, 19, 724.

[12] A. M. Salvi, K. A. DeMali, Curr. Opin. Cell Biol. 2018, 54, 114.

[13] P. Romani, I. Brian, G. Santinon, A. Pocaterra, M. Audano, S. Pedretti, S. Mathieu, M. Forcato, S. Bicciato, J.-B. Manneville, Nat. Cell Biol. 2019, 338.

[14] E. Bartoláksuki, J. Imsirovic, H. Parameswaran, T. J. Wellman, N. Martinez, P. G. Allen, U. Frey, B. Suki, Nat. Mater. 2015, 14, 1049.

[15] M. R. Zanotelli, A. Rahman-Zaman, J. A. VanderBurgh, P. V. Taufalele, A. Jain, D. Erickson, F. Bordeleau, C. A. Reinhart-King, Nat. Comm. 2019, 10, 1.

[16] H. Zhao, T. Li, K. Wang, F. Zhao, J. Chen, G. Xu, J. Zhao, T. Li, L. Chen, L. Li, Q. Xia, T. Zhou, H. Y. Li, A. L. Li, T. Finkel, X. M. Zhang, X. Pan, Nat. Cell Biol. 2019, DOI: 10.1038/s41556-019-0296-3476.

[17] B. B. Kahn, A. Thierry, C. David, D. G. Hardie, Cell Metabol. 2005, 1, 15.

[18] A. Avivar-Valderas, E. Bobrovnikova-Marjon, J. A. Diehl, N. Bardeesy, J. Debnath, J. A. Aguirre-Ghiso, (2013). Oncogene 2013, 32, 4932.

[19] D. G. Hardie, D. Carling, M. Carlson, Annu. Rev. Biochem. 1998, 67, 821.

[20] M. Jung-Soon, M. Zhipeng, K. Young Chul, P. Hyun Woo, H. Carsten Gram, K. Soohyun, L. Dae-Sik, G. Kun-Liang, Nat. Cell Biol. 2015, 17, 500.

[21] D. G. Hardie, F. A. Ross, S. A. Hawley, Nat. Rev. Mol. Cell Biol. 2012, 13, 251.

[22] S. C. J. Helle, Q. Feng, M. J. Aebersold, L. Hirt, R. R. Grüter, A. Vahid, A. Sirianni, S. Mostowy, J. G. Snedeker, A. Šarić, Elife 2017, 6, e30292.

[23] A. S. Rambold, B. Kostelecky, N. Elia, J. Lippincott-Schwartz, Proc. Natl. Acad. Sci. 2011, 108, 10190.

[24] E. Q. Toyama, S. Herzig, J. Courchet, L. T. Jr, O. C. Losón, K. Hellberg, N. P. Young, H. Chen, F. Polleux, D. C. Chan, Science 2016, 351, 275.

[25] S. Herzig, R. J. Shaw, Nat. Rev. Mol. Cell Biol. 2018, 19, 121.

[26] K. Inoki, T. Zhu, K. L. Guan, Cell 2003, 115, 577.

[27] K. R. Laderoute, K. Amin, J. M. Calaoagan, M. Knapp, T. Le, J. Orduna, M. Foretz, B. Viollet, Mol. Cell. Biol. 2006, 26, 5336.

[28] J. T. Connelly, J. E. Gautrot, B. Trappmann, D. W.-M. Tan, G. Donati, W. T. Huck, F. M. Watt, Nat. Cell Biol. 2010, 12, 711.

[29] M. Bao, J. Xie, A. Piruska, W. T. Huck, Nat. Comm. 2017, 8, 1962.

[30] W. J. Koopman, F. Distelmaier, J. J. Esseling, J. A. Smeitink, P. H. Willems, Methods 2008, 46, 304.

[31] M. Deschepper, M. Manassero, K. Oudina, J. Paquet, L. E. Monfoulet, M. Bensidhoum, H. Petite, Stem Cells 2013, 31, 526.

[32] D. Sirio, M. Leonardo, A. Mariaceleste, E. Elena, G. Stefano, C. Michelangelo, Z. Francesca, L. D. Jimmy, F. Mattia, B. Silvio, Nature 2011, 474, 179.

[33] M. Khacho, A. Clark, D. S. Svoboda, J. Azzi, J. G. Maclaurin, C. Meghaizel, H. Sesaki, D. C. Lagace, M. Germain, M. E. Harper, Cell Stem Cell 2016, 19, 232

[34] Z. Meng, Y. Qiu, K. C. Lin, A. Kumar, J. K. Placone, C. Fang, K. C. Wang, S. Lu, M. Pan, A. W. Hong, T. Moroishi, M. Luo, S. W. Plouffe, Y. Diao, Z. Ye, H. W. Park, X. Wang, F. X. Yu, S. Chien, C. Y. Wang, B. Ren, A. J. Engler, K. L. Guan, Nature 2018, 560, 655.

[35] L. Valon, A. Marín-Llauradó, T. Wyatt, G. Charras, X. Trepat, Nat. Comm. 2017, 8, 14396.

[36] K. Clark, M. Langeslag, C. G. Figdor, F. N. van Leeuwen, Trends Cell Biol. 2007, 17, 178.

[37] Y. Chun, M. W. Tibbitt, B. Lena, K. S. Anseth, Nat. Mater. 2014, 13, 645.

[38] Z. Zhuoran, Z. Ming, X. Guozhi, R. T. Franceschi, Mol. Ther. 2005, 12, 247.

[39] J. D. Humphrey, E. R. Dufresne, M. A. Schwartz, Nat. Rev. Mol. Cell Biol. 2014, 15, 802.

[40] L. C. Kelley, Q. Y. Chi, R. Cáceres R, E. Hastie, A. J. Schindler, Y. Jiang, D. Q. Matus, J. Plastino, D. R. Sherwood, Dev. Cell 2019, 48, 313.

[41] T. Shiraishi, J. E. Verdone, J. Huang, U. D. Kahlert, J. R. Hernandez, G. Torga, J. C. Zarif, T. Epstein, R. Gatenby, A. M. Cartney, J. H. Elisseeff, S. M. Mooney, K. J. Pienta, Oncotarget 2015, 6, 130.

[42] E. J. Mah, A. E. Lefebvre, G. E. McGahey, A. F. Yee, M. A. Digman, Sci. Rep. 2018, 8, 1.

[43] M. R. Zanotelli, Z. E. Goldblatt, J. P. Miller, F. Bordeleau, J. Li, J. A. VanderBurgh, M. C. Lampi, M. R. King, C. A. Reinhart-King, Mol. Bio. Cell, 2018, 29, 1.

[44] S. Tojkander, K. Ciuba, P. Lappalainen, Cell Rep. 2018, 24, 11.

[45] S. Tojkander, G. Gateva, A. Husain, R. Krishnan, P. Lappalainen, Elife 2015, 4, e06126.

[46] B. Cunniff, A. J. McKenzie, N. H. Heintz, A. K. Howe, Mol. Biol. Cell, 2016, 27, 2662.

[47] M. H. Schuler, A. Lewandowska, G. D. Caprio, W. Skillern, S. Upadhyayula, T. Kirchhausen, J. M. Shaw, B. Cunniff, Mol. Biol. Cell, 2017, 8, 2159.

[48] J. S. Park, C. J. Burckhardt, R. Lazcano, L. M. Solis, T. Isogai, L. Q. Li, C. S. Chen, B. N. Gao, J. D. Minna, R. Bachoo, R. J. DeBerardinis, G. Danuser, Nauture, 2020, 578, 621

