## supplementary information for "Energy Expenditure during Cell Spreading Regulates the Stem Cells Responses to Matrix Stiffness"

### Supplementary Figures

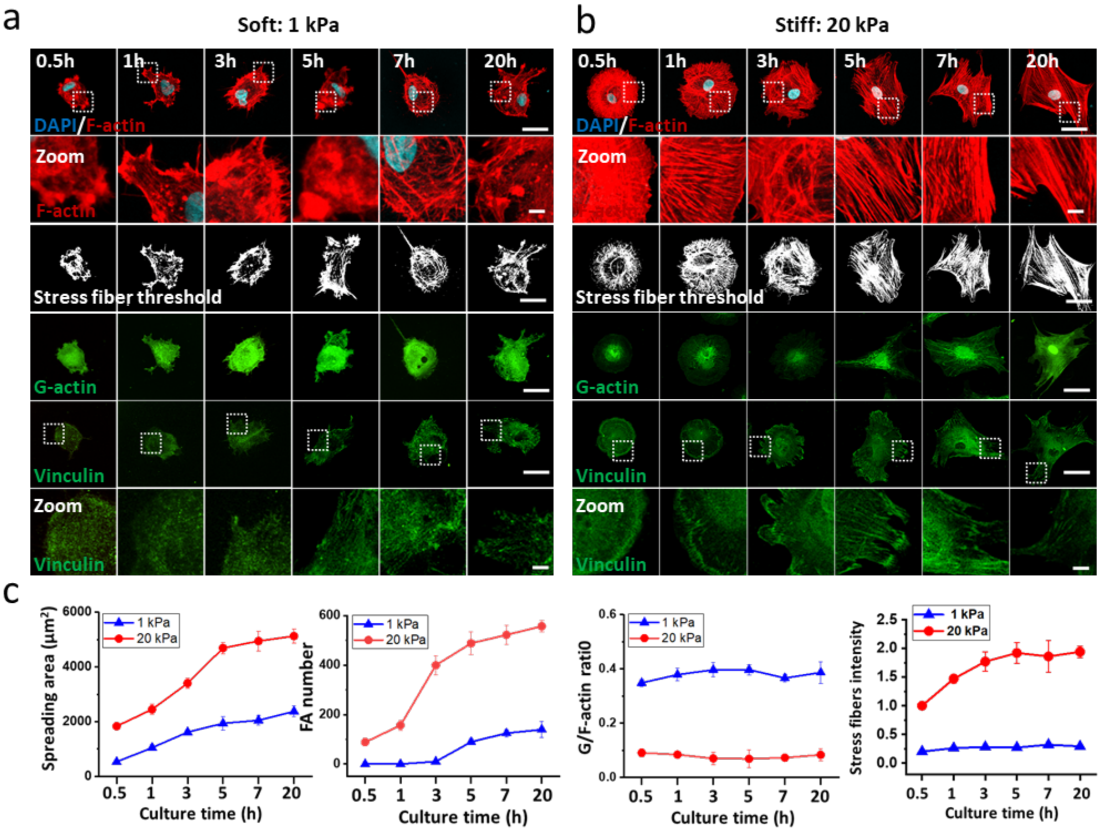


**Figure S1. hMSCs display different F-actin organization and FA formation over time on PAAm gels with different stiffness.** (A, B) Confocal images of hMSCs cultured onto (A) soft (1 kPa) and (B) stiff (20 kPa) PAAm gels showing G-actin (DNaseI-labeled staining: green) and FAs (vinculin staining: green) at different time points. F-actin was stained with phalloidin (red) and nuclei were counterstained with DAPI (blue). Scale bars, 50 μm and 10 μm (zoom). (C) Quantification of spreading area,  FA number, G-actin to F-actin ratio and stress fibers intensity over time on PAAm gels with different stiffness (n>75 cells from three independent experiments).

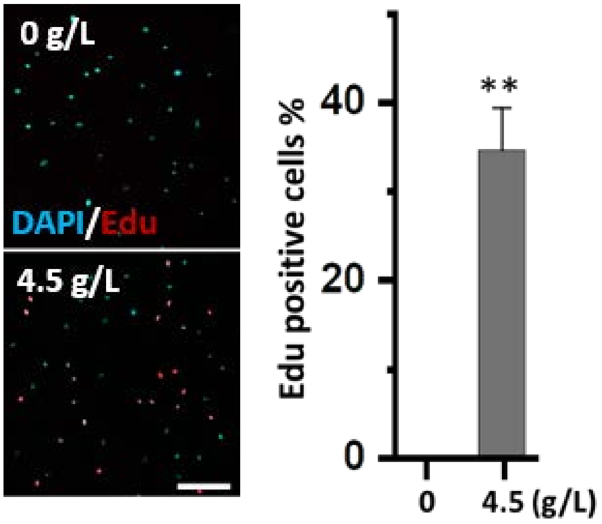


**Figure S2. Energy starvation greatly reduces hMSCs proliferation on stiff PAAm gels.** The confocal images showed the merged view of EdU (red) and DAPI (blue) at the presence (bottom) or absence (top) of glucose. Graph: quantification of Edu positive cells (n>18 representative images from three independent experiments). Scale bars, 200 μm.

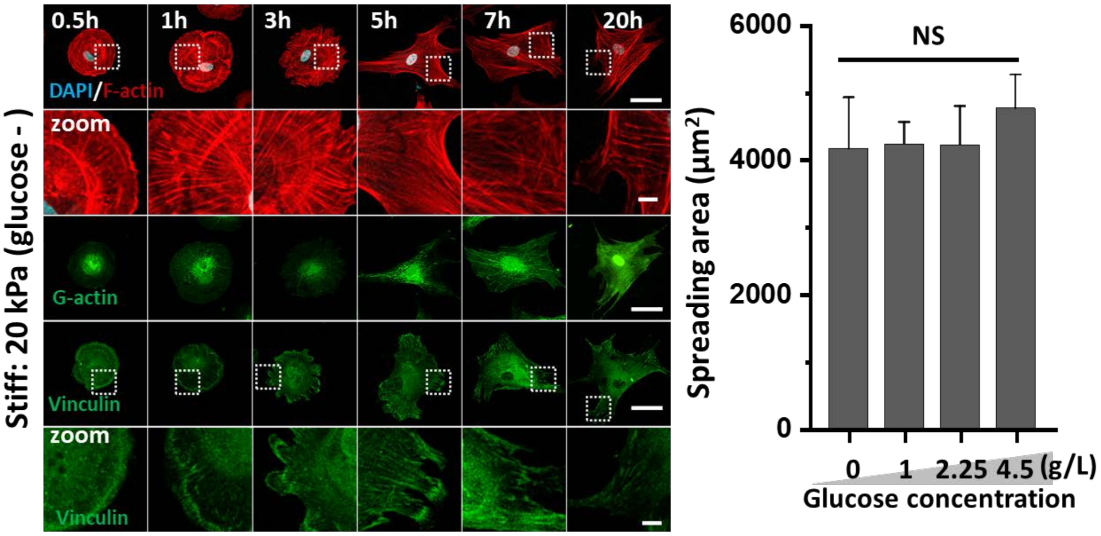


**Figure S3. Glucose starvation limits FA formation of hMSCs cultured on stiff PAAm gels, while having no effect on the spreading area 20 h after seeding.** Confocal images of hMSCs cultured with glucose starvation on stiff PAAm gel showing G-actin (DNaseI-labeled staining: green) and FAs (vinculin staining: green) at different time points. F-actin was stained with phalloidin (red) and nuclei were counterstained with DAPI (blue). Scale bars, 50 μm. Graph: quantification of spreading area under different glucose concentration after 20 h culture of cells. (n>40 cells from three independent experiments).

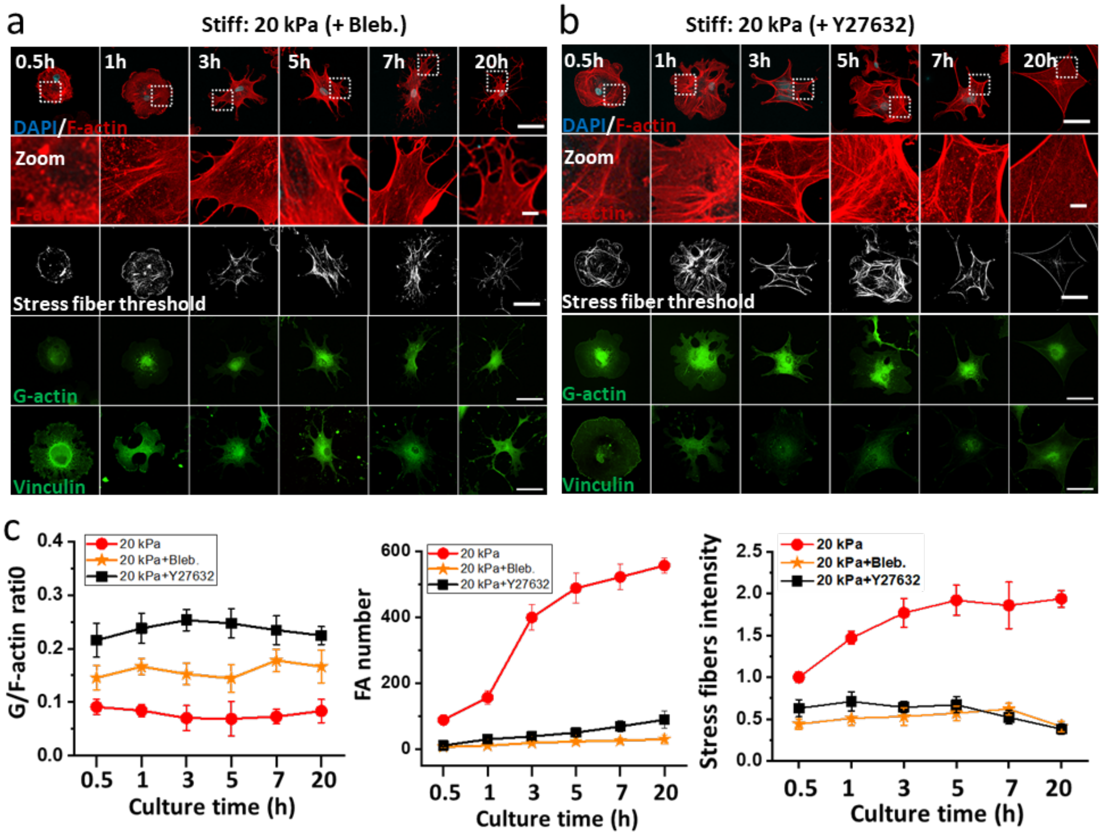


**Figure S4. F-actin organization and FA number of hMSCs is influenced by inhibition of cellular contractility.** (A, B) Confocal images of hMSCs cultured onto stiff substrate treated with (A) myosin inhibitor (50 μM Bleb.) or (B) Rock inhibitor (50 μM Y27632) showing G-actin (DNaseI-labeled staining: green), FAs (vinculin staining: green) and stress fibers at different time points. F-actin was stained with phalloidin (red) and nuclei were counterstained with DAPI (blue) (C) Quantification of G/F actin ratio FA number and stress fibers intensity of hMSCs treated with contractility inhibition on stiff (20 kPa) substrates. (n>75 cells from three independent experiments).

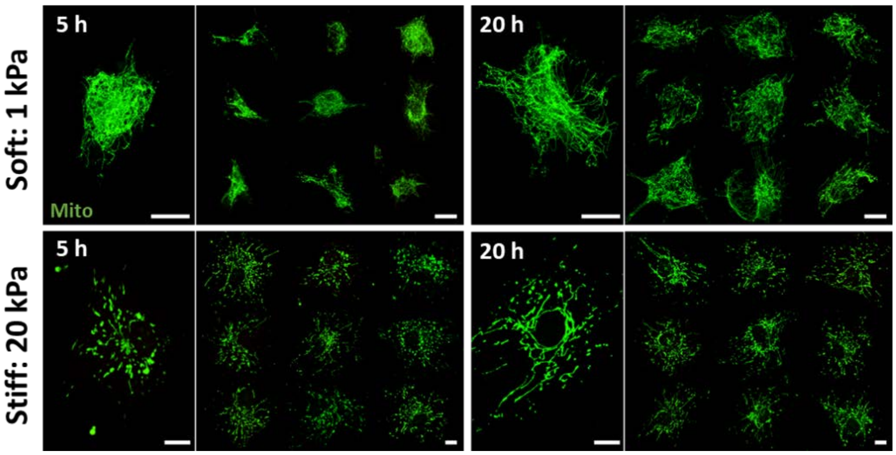


**Figure S5 More representative images for mitochondrial morphologies of hMSCs cultured on soft or stiff PAAm gels at 5 or 20 hours after seeding**. Mitochondria were stained with Mito tracker (green). Scale bars, 25μm.

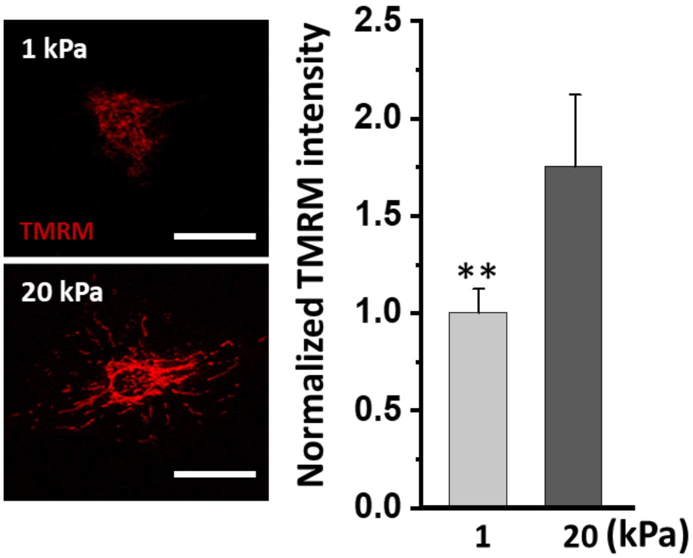


**Figure S6. Mechanical properties of the substrate affect mitochondrial TMRM accumulation linked to mitochondrial membrane potential.** hMSCs cultured on soft (1 kPa) or stiff substrates (20 kPa) displayed different mitochondrial TMRM intensities. Scale bars, 50µm. Graph: quantification of average mitochondrial pixel intensity (n>60 cells from three independent experiments).

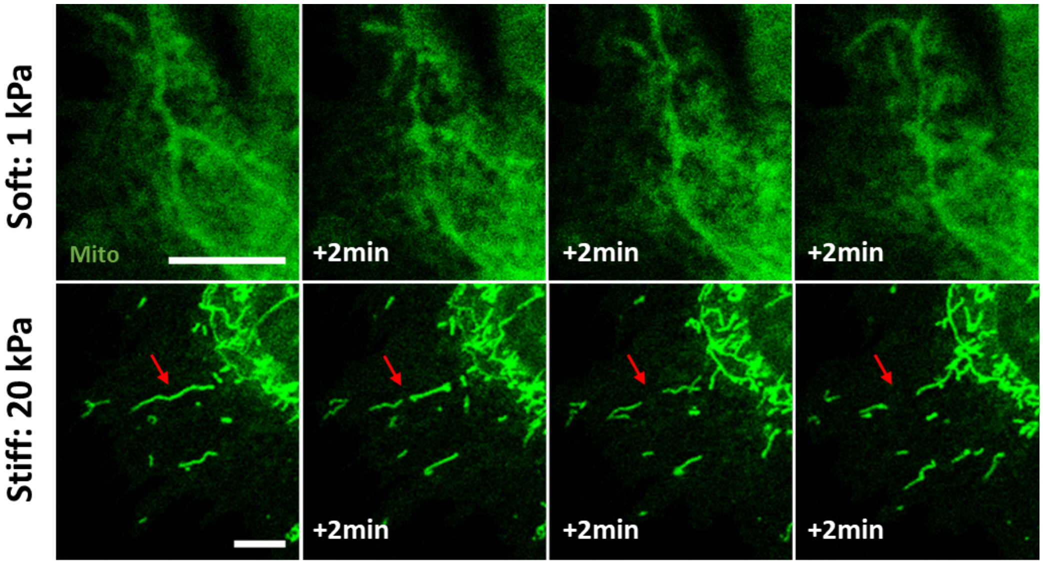


**Figure S7. Mitochondrial fission event on a stiff substrate.** Time-lapse images of hMSCs cultured on PAAm gels of different stiffness stained with Mito tracker. The red arrow highlights a mitochondrial fission on the stiff substrate. Scale bars, 10 µm

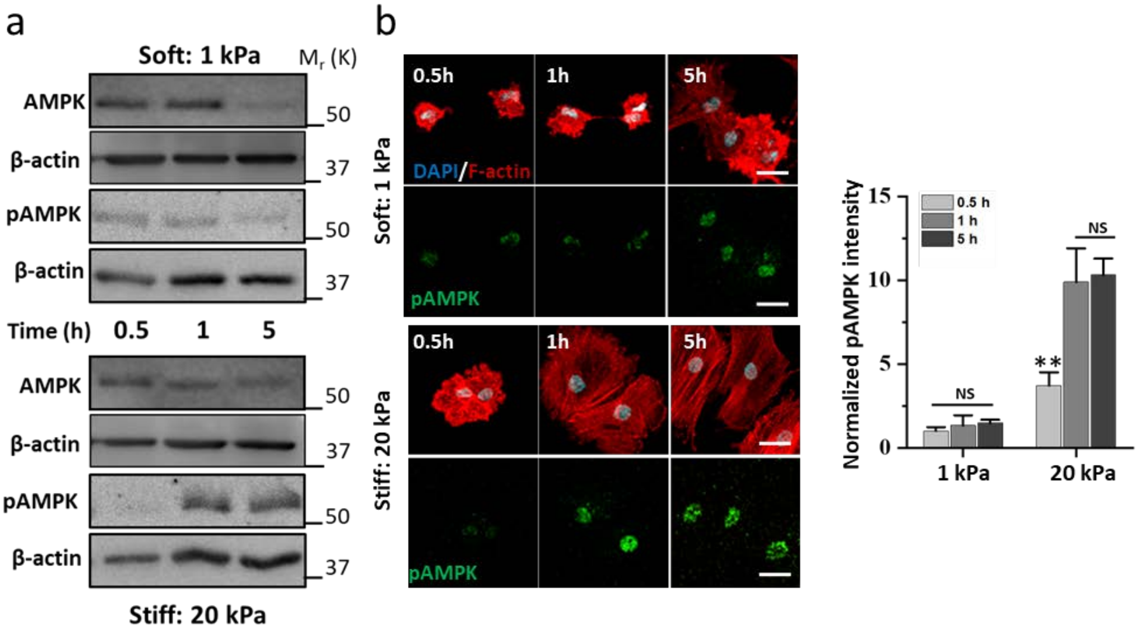


**Figure S8. AMPK activation of hMSCs starts from 1 h on stiff substrate.** (A) Western blots of AMPK and pAMPK of cells at different time points after seeding. (B) Confocal images of hMSCs stained with antibodies against pAMPK (green) and counterstained with DAPI (blue) and actin (red) at different time points after seeding. Scale bars, 50 µm. Graph: quantification of fluorescence intensity of nuclear pAMPK in single cells cultured on different PAAm gels at different time points (n>10 representative images from two independent experiments).

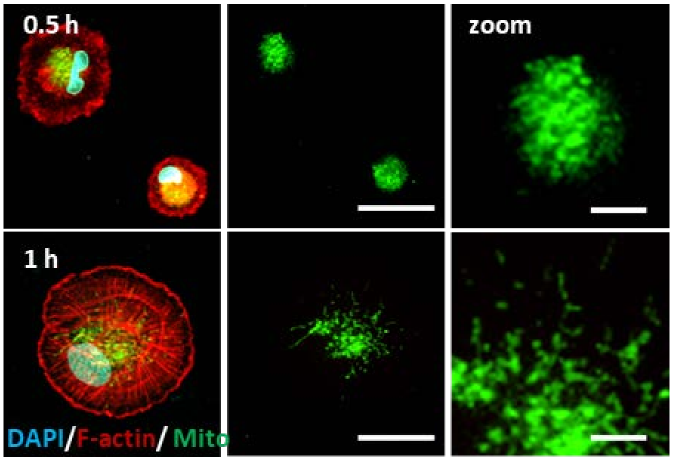


**Figure S9. Mitochondria of hMSCs seeded on a stiff substrate display fragmentation from 1 h onward.** Mitochondrial morphologies on stiff PAAm gels at 0.5 or 1 h after seeding. Mitochondria are stained with 100 nM MitoTracker (green). Scale bars, 25 µm and 5 µm in zoomed images. Images are representative of at least three independent experiments with similar results.

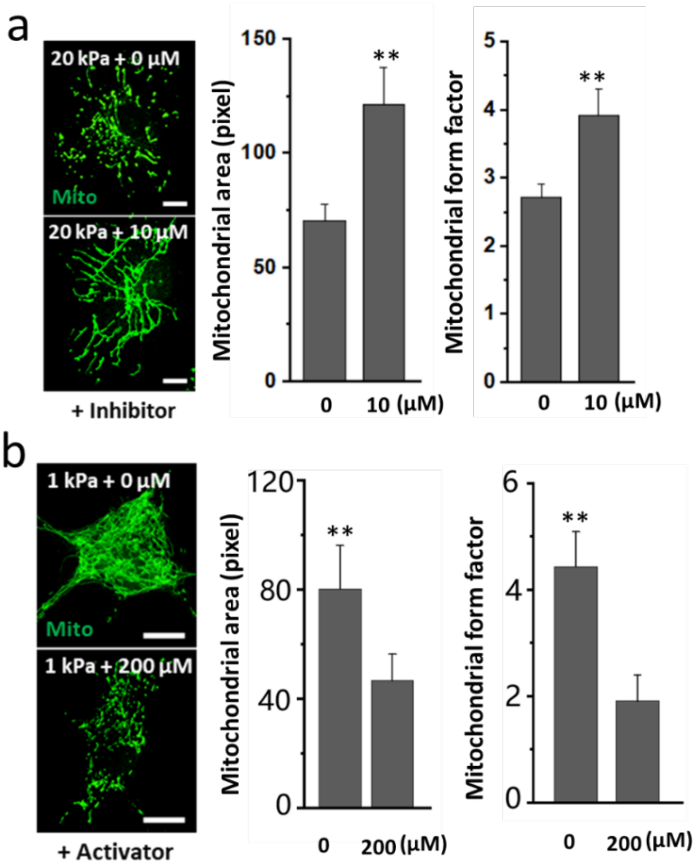


**Figure S10. AMPK inhibition or activation alters mitochondrial morphology.** Quantification of mitochondrial area and form factor of hMSCs treated by (A) AMPK inhibitor (Compound C) or (B) activator (A-769662) using the measurement plugin of Fiji software. Scale bars, 20 μm (a) and 10 μm (b), n>15 cells from two independent experiment.

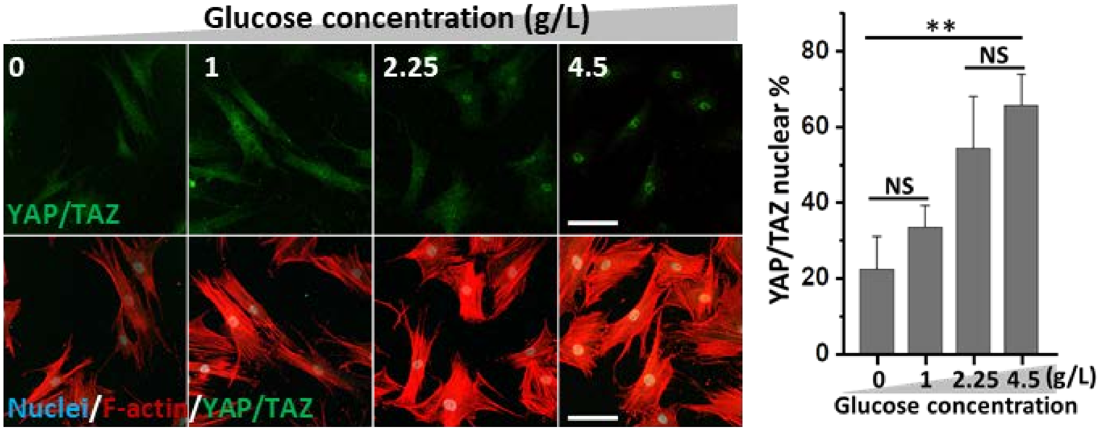


**Figure S11. Energy starvation inhibits YAP/TAZ nuclear localization on stiff substrate.** Images of YAP/TAZ localization in hMSCs cultured in the medium with different glucose concentration. Scale bars, 100 μm. Graph: Quantification of YAP/TAZ nuclear localization under different culture conditions (n>40 cells from three independent experiments).

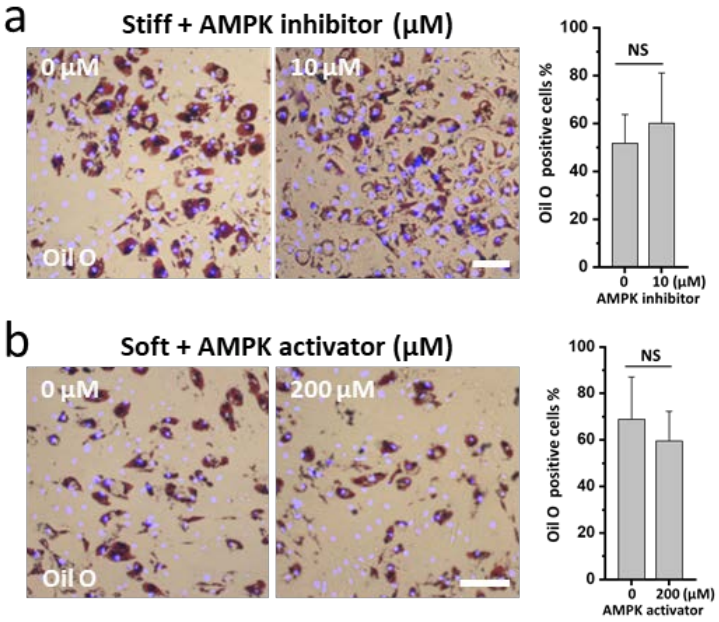


**Figure S12. AMPK activation or inhibition does not affect adipogenic differentiation of hMSCs.** Oil O staining showed adipogenic differentiation of hMSCs seeded on different substrates. Oil O (bright field) expression of hMSCs treated with AMPK inhibition (10 µM Compound C) on stiff substrates (A) or AMPK activation (200 µM A-769662) on soft substrates (B). Nuclei were counterstained with DAPI (blue). Scale bars, 150 µm. Graph: quantification Oil O positive cells after 10 day (n>6 representative images from two independent experiments) at different conditions.

### Movie S1. Mitochondrial dynamics of hMSCs cultured on soft (1 kPa) or stiff (20 kPa) PAAm gels

Mitochondria were imaged every 2 min from 3 h after seeding to track the mitochondrial dynamics. Scale bars, 50 µm
